# Single Cell Transcriptomics and Flow Cytometry Reveal Disease-associated Fibroblast Subsets in Rheumatoid Arthritis

**DOI:** 10.1101/126193

**Authors:** Fumitaka Mizoguchi, Kamil Slowikowski, Jennifer L Marshall, Kevin Wei, Deepak A Rao, Sook Kyung Chang, Hung N Nguyen, Erika H Noss, Jason D Turner, Brandon E Earp, Philip E Blazar, John Wright, Barry P Simmons, Laura T Donlin, George D Kalliolias, Susan M Goodman, Vivian P Bykerk, Lionel B Ivashkiv, James A Lederer, Nir Hacohen, Peter A Nigrovic, Andrew Filer, Christopher D Buckley, Soumya Raychaudhuri, Michael B Brenner

## Abstract

Fibroblasts mediate normal tissue matrix remodeling, but they can cause fibrosis or tissue destruction following chronic inflammation. In rheumatoid arthritis (RA), synovial fibroblasts expand, degrade cartilage, and drive joint inflammation. Little is known about fibroblast heterogeneity or if aberrations in fibroblast subsets relate to disease pathology. Here, we used an integrative strategy, including bulk transcriptomics on targeted subpopulations and unbiased single-cell transcriptomics, to analyze fibroblasts from synovial tissues. We identify 7 phenotypic fibroblast subsets with distinct surface protein phenotypes, and these collapsed into 3 subsets based on transcriptomics data. One subset expressing PDPN, THY1, but lacking CD34 was 3-fold expanded in RA relative to osteoarthritis (P=0.007); most of these cells expressed CDH11. The subsets were found to differ in expression of cytokines and matrix metalloproteinases, localization in synovial microanatomy, and in response to TNF. Our approach provides a template to identify pathogenic stromal cellular subsets in complex diseases.

## Main Text

Fibroblasts are key mediators of end organ pathology and inflammation in many diseases. Although they mediate normal matrix deposition and inflammation in wound healing, chronically activated fibroblasts can differentiate into myofibroblasts that lay down collagen and are responsible for fibrosis in lung, liver, gut, skin, and other tissues (*1*). Conversely, chronically activated fibroblasts are responsible for excessive matrix degradation that destroys cartilage and causes permanent joint damage in rheumatoid arthritis (*2–4*). Moreover, recent studies have emphasized the role of fibroblasts as stromal cells that regulate immune responses in lymph nodes and tumor stroma (*5, 6*). Unlike hematopoietic cell types that are known to be comprised of a variety of functionally different cellular types and subsets, fibroblasts have generally been considered to have little heterogeneity: functionally distinct subpopulations have yet to be clearly defined.

Recent advances in high-throughput technologies enable investigators to query complex diseases in humans in new ways. For example, global transcriptomic analysis revealed distinct activation states and cellular subsets of immune cells (*7*). These approaches offer an opportunity to determine how stromal cells mediate various types of local tissue pathology. Transcriptomics of small numbers of cells, and even single cells, from human pathological samples can advance the understanding of tissue dynamics in disease. For example, single cell RNA-seq has revealed heterogeneity of tumor cells and identified a mechanism for drug resistance in cancer (*8, 9*).

Here, we focus on mesenchymal cell heterogeneity in RA, a complex autoimmune disease affecting up to 1% of the world's population (*10*). In RA, the synovium changes dramatically: the thin membrane encapsulating the joint becomes an inflamed, hyperplastic, and invasive tissue mass that causes joint destruction (*4*). Synovial fibroblasts secrete inflammatory cytokines and chemokines, invade and degrade cartilage, and stimulate osteoclasts that cause bone erosion (*2, 4*).

We hypothesized that these different functions might be carried out by distinct cellular subsets of fibroblasts, analogous to functionally distinct subsets of leukocytes. If fibroblast subsets exist, we thought that altered proportions of fibroblast subsets might underlie pathological observations in tissues (*11*). Since there are no approved drugs that directly target fibroblasts, identifying pathogenic fibroblast subsets may reveal therapeutic targets broadly applicable across a range of diseases. In RA, targeting fibroblast subsets might complement anti-inflammatory therapies that target leukocytes and their cytokines (*4*).

## Results

### Fibroblasts in synovial tissue have distinct surface markers

To examine the heterogeneity of fibroblasts in joint tissue, we isolated cells by enzymatic digestion of synovial specimens from joint replacement surgery. We compared RA samples to osteoarthritis (OA) samples. OA samples have been widely used as controls or comparators in the studies of RA, since OA is a degenerative joint disease while RA is an autoimmune disease associated with synovial hypertrophy and inflammation. We first examined protein expression of a variety of surface markers on fibroblasts including podoplanin (PDPN), cadherin 11 (CDH11), THY1 (CD90), and CD34 by flow cytometry (see **Supplementary Methods**). To isolate fibroblasts, we first gated PTPRC(CD45)^−^GYPA(CD235a)^−^PECAM1(CD31)^−^ cells to remove hematopoietic lineage cells, red blood cells (RBCs), and endothelial cells, and further removed MCAM(CD146)^+^ pericytes. The remaining cells had high PDPN expression, consistent with fibroblasts in the synovium (**Fig 1A**). We identified two major populations based on the expression of CD34. In 42 donors (26 OA and 16 RA), we observed medians of 34.7% CD34^+^ and 54.7% CD34^−^ cells (**Fig 1A, Table S1**). CD34^−^ and CD34^+^ fibroblasts could further be divided into 4 and 3 populations, respectively, based on THY1 and CDH11 expression. However, we remained uncertain about which of these divisions were functionally meaningful. This result prompted us to further investigate if the gated fibroblast populations might have different gene expression profiles and effector functions.

**Fig. 1.**
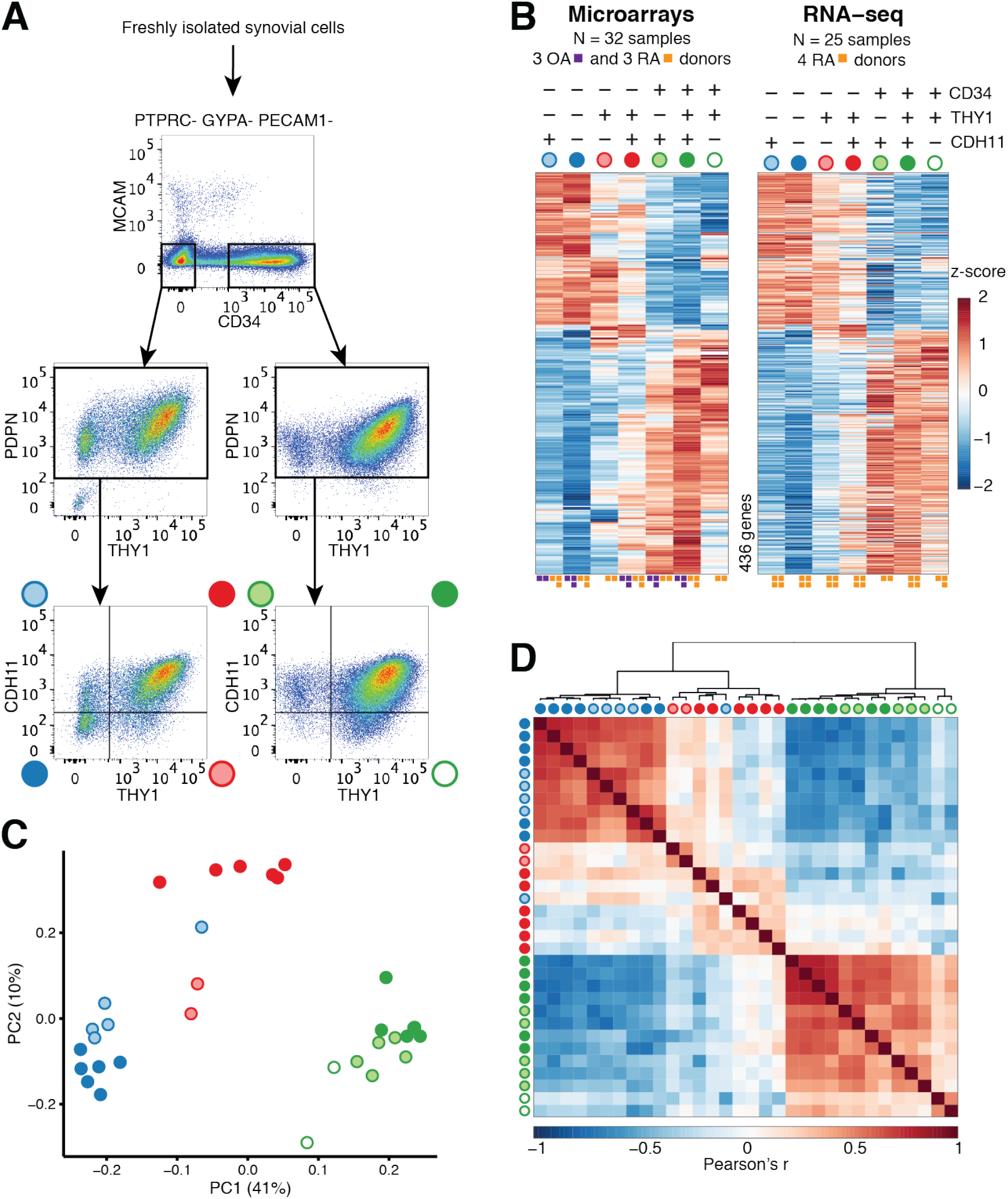
Fibroblast subsets have distinct protein and mRNA expression. (**A**) Gating strategy for synovial fibroblasts with heterogeneous expression of surface proteins. (**B**) Analysis of variance (ANOVA) reveals 436 genes with significant (1% FDR) variation across 7 gated populations that are measured and statistically significant in both microarray and RNA-seq datasets. Squares beneath each column indicate number of samples for that phenotypic subset. (**C**) Principal components analysis (PCA) with 2,986 genes (1% FDR, ANOVA) in microarray data separates the 32 microarray samples into 3 subsets: CD34^−^THY1^−^, CD34^−^THY1^+^, and CD34^+^. (**D**) Pairwise Pearson correlation of microarrays also suggests 3 major subsets of fibroblasts.

### Fibroblast subpopulations have three distinct gene expression signatures

We used two complementary strategies to investigate synovial fibroblast populations obtained after tissue disaggregation: (1) fluorescence sorting using a set of candidate protein markers followed by bulk transcriptomics of gated populations and (2) unbiased single cell transcriptomics without gating.

First, we assayed each of the 7 gated populations from 3 OA and 3 RA donors with the Affymetrix HuGene 2.0 ST microarray, using Robust Multichip Average to normalize 53,617 probesets and 20,452 genes. As expected, we observed that all of the samples expressed genes typically expressed in fibroblasts and lacked expression of other lineage specific genes (**Fig. S1A**). However, after controlling for variation between donors, we observed 2,986 genes with significant variation across the 7 distinct populations gated based on surface protein marker expression (FDR<1%, analysis of variance (ANOVA)), suggesting notable transcriptional differences across these putative subsets. To validate these microarray findings, we applied RNA-seq to the same 7 subsets from 4 independent RA donors. We prepared libraries with Smart-Seq2, sequenced to an average depth of 5.6M fragments per sample, and quantified expression for 19,532 genes. These samples were also enriched with fibroblast lineage genes (**Fig. S1B**). Those genes that were significantly differentially expressed in the microarray experiment and that overlapped with the RNA-seq experiment (n=2,659 genes) had similar expression profiles across the 7 subsets (**Fig. S2**). In total, 436 genes were measured on both platforms and had significant variation across the 7 putative subpopulations in both experiments (1% FDR, ANOVA) (**Fig. 1B**). The concordance of these results suggests that the gene expression differences reflect biological variation rather than technical or stochastic artifacts. We reasoned that these expression profiles could serve as proxies for molecular functions to define putative cellular subsets with distinct biological roles.

Principal component analysis revealed that 7 phenotypic populations fall into 3 distinct major subsets: CD34^−^THY1^−^, CD34^−^THY1^+^, and CD34^+^. Principal component 1 (PC1) clearly separates CD34^−^ and CD34^+^ samples and PC2 separates the CD34^−^ samples that are THY1^+^ and CDH11^+^ (**Fig. 1C**). The 3 subsets were also clearly apparent by hierarchical clustering on pairwise Pearson correlations (**Fig. 1D**). CD34^−^ and CD34^+^ samples were positively correlated within subsets and negatively correlated between subsets. CD34^−^THY1^+^ sample correlations were less consistent between the 3 subsets and within this subset, indicating that this subset may be more heterogeneous than the other two (**Fig. 1D**). We decided to group CD34^−^THY1^−^CDH11^−^ and CD34^−^ THY1^−^CDH11^+^ samples because they had similar gene expression profiles overall.

### Single cell RNA-seq confirmed the presence of three major fibroblast subsets

Next, since our observation from gated fibroblast populations may be potentially biased by our a priori selection of surface markers, we performed single cell mRNA sequencing to obtain an unbiased characterization of transcriptional heterogeneity in fibroblasts. We depleted hematopoietic cells, RBCs, and endothelial cells, and selected cells positive for PDPN (PTPRC^−^ GYPA^−^ PECAM1^−^ PDPN^+^) (**Fig. S1C**). We used Illumina Smart-Seq2 to prepare single cell libraries from fibroblasts from 4 additional donors, 2 RA and 2 OA; we sequenced to an average depth of 5.4M fragments per cell, and detected an average of 8,842 genes per cell with at least 1 transcript per million (TPM) (**Fig. S3**). We selected 337 cells with at least 5,000 detected genes, and then used Single Cell Differential Expression (SCDE) to estimate error models for each cell, normalize expression values, remove aspects of variation due to library complexity, and correct for batch effects across donors (*12*). We measured protein levels on the single cells by flow cytometry at the time of single cell sorting for comparison with the single cell transcriptomic analyses. Remarkably, unbiased subsets (defined by hierarchical clustering of single cells with 23 genes that show high mean and variance across cells) were concordant with the three subsets defined by protein levels of CD34 and THY1 surface markers (permutation P<10^−4^) (**Fig. 2A, B**).

**Fig. 2.**
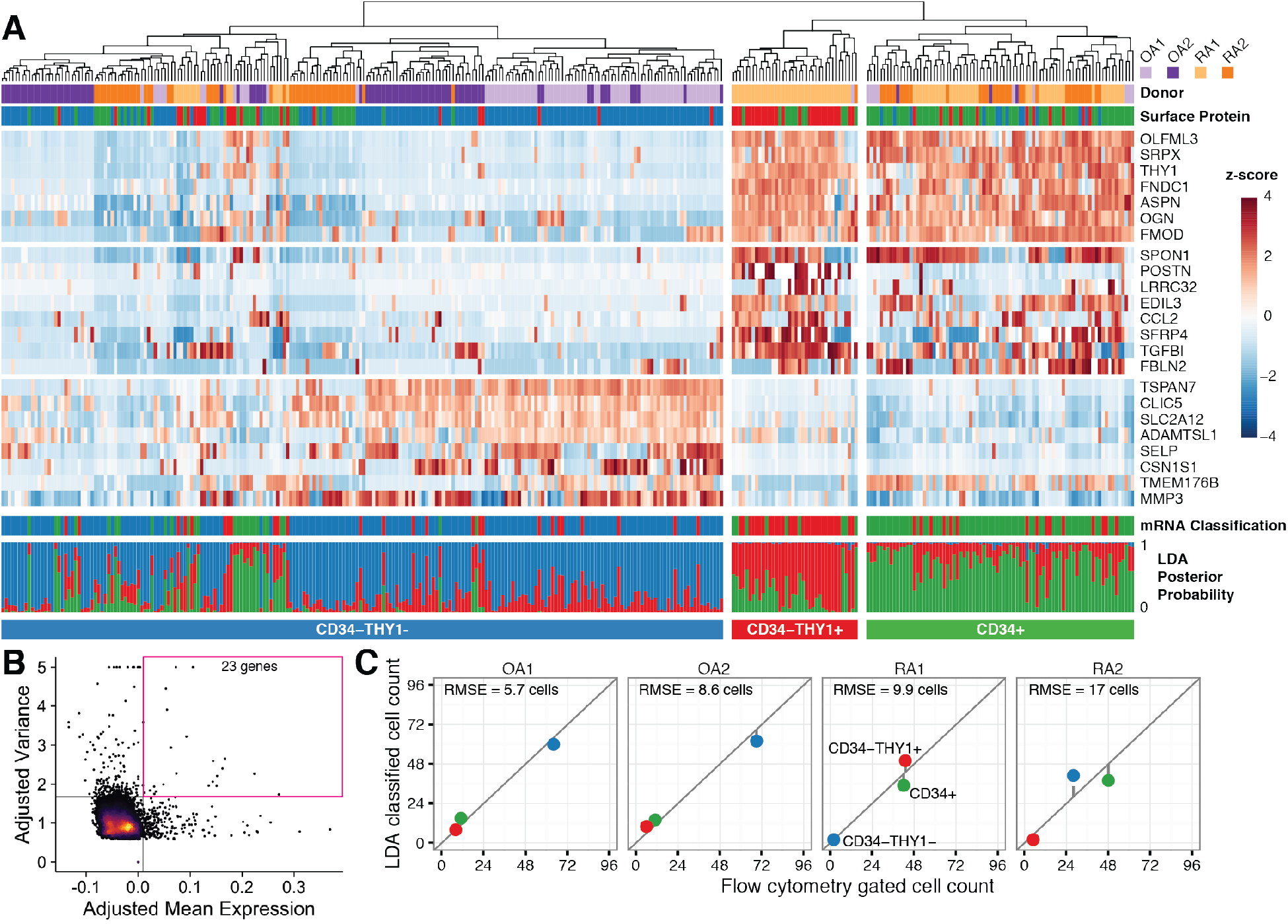
Single-cell RNA-seq validates findings in bulk assays. (**A**) Hierarchical clustering is concordant with our surface protein gating. Bottom tracks show linear discriminant analysis (LDA) classification by mRNA expression of 968 genes and posterior probabilities. (**B**) For the heatmap, we selected 23 genes with greatest 1% mean and greatest 1% variance in expression levels. (**C**) Number of cells in each subset as determined by surface protein levels (x-axis) and by LDA classification (y-axis). Root mean squared error (RMSE).

Next, we assessed whether the bulk and single cell RNA-seq profiles were marking similar cellular subpopulations. We trained a linear discriminant analysis (LDA) classifier on 968 genes in the bulk RNA-seq data and predicted the classes of single cells (see **Supplementary methods**). The LDA classifier produced a probability of belonging to each class (CD34^−^THY1^−^, CD34^−^THY1^+^, and CD34^+^). The classifications were confident (median probability 0.8) and consistent with the three subsets (**Fig. 2A**). We compared mRNA classification with flow cytometric protein marker data recorded at the time of unbiased single cell sorting and found concordant cell identities and proportions (**Fig. 2C, S4**). These single-cell RNA-seq results provide an unbiased and independent validation of the 3 major subsets defined with bulk transcriptomics of samples gated by protein surface markers.

### Fibroblast subsets localize to specific regions of lymphoid structures in the synovium

Next, we characterized the anatomical localization of fibroblast subsets in synovial tissue and assessed differences between OA and RA (**Fig. 3A, B**). CD34^−^ THY1^+^ fibroblasts in RA samples appeared as expanded masses of cells in the deeper sublining layer of the synovium surrounding blood vessels, especially near accumulations of lymphocytes. In OA samples, the CD34^−^THY1^+^ fibroblasts comprise a thin layer with fewer cells surrounding blood vessels than in RA samples. CD34^+^ fibroblasts were observed in both superficial lining and deeper sublining areas of the synovium. CD34^−^THY1^−^ fibroblasts were mostly observed in lining area.

**Fig. 3.**
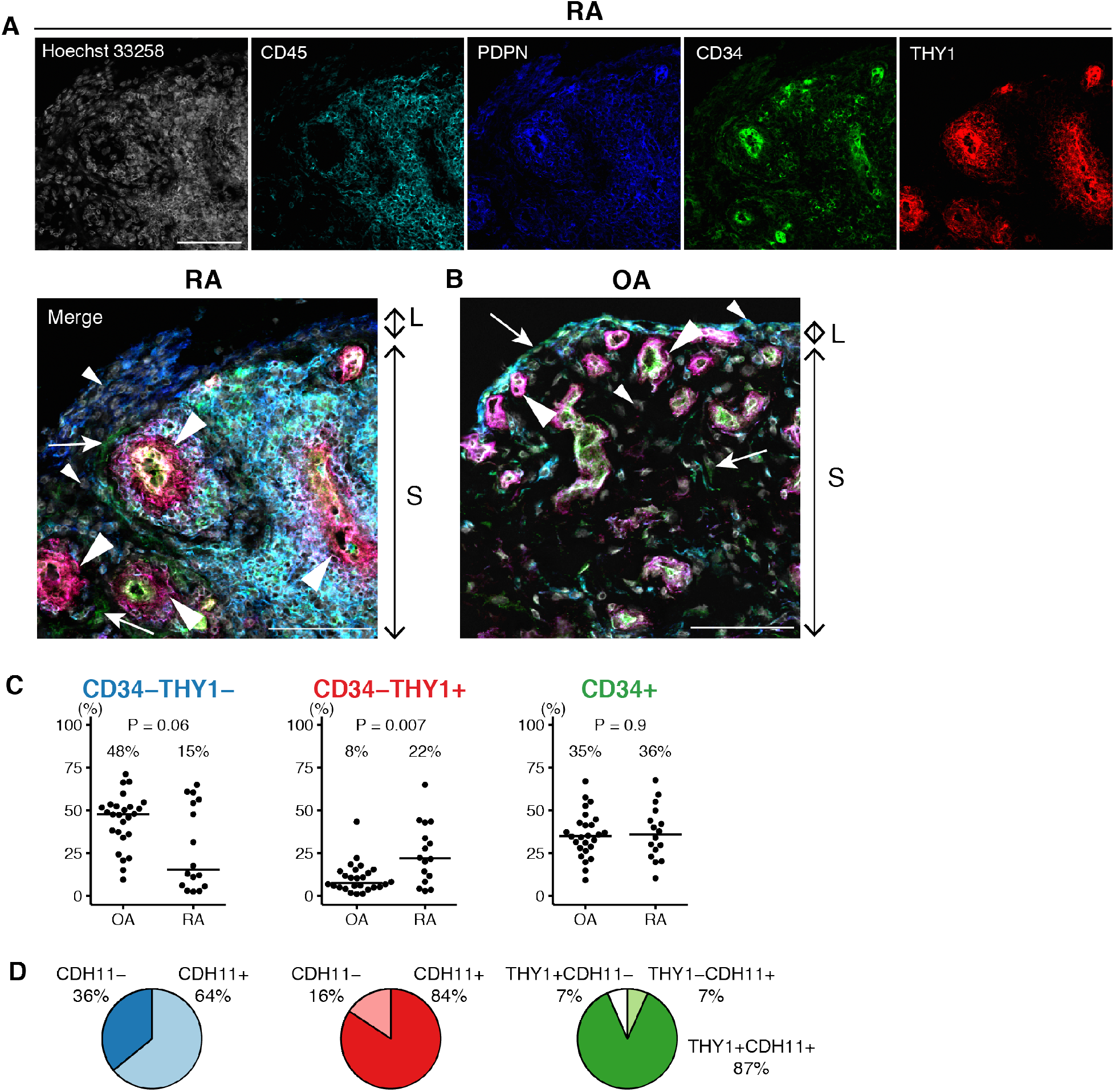
The anatomical localization and the proportion of fibroblast subsets in synovial tissue are different between OA and RA. (**A, B**) Anatomical localization of fibroblast subsets in RA and OA synovial tissue. Hoechst 33258: White, CD45: Cyan, PDPN: Blue, CD34: Green, THY1: Red. Small arrow heads: CD34^−^THY1^−^ fibroblasts. Big arrow heads: CD34^−^THY1^+^ fibroblasts. Arrows: CD34^+^ fibroblasts. L: Lining area. S: Sublining area. Scale = 100 μm. (**C**) Proportions of fibroblast subsets in synovial tissue in OA (n=26) and RA (n=16) evaluated by flow cytometry. (**D**) Proportions of CDH11^+^ cells in CD34^−^THY1^−^ fibroblasts, CD34^−^THY1^+^ fibroblasts and CD34+ fibroblasts.

### Fibroblast subset proportions are different between RA and OA

We hypothesized that if pathological fibroblast subsets exist, then some subsets might be more or less abundant in RA relative to OA synovial tissues. Indeed, proportions of fibroblast subsets defined by flow cytometry with protein surface markers were different between RA synovial tissue (n=16) and OA (n=26) (**Fig. 3C, Table S1**). The proportion of CD34^−^THY1^+^ fibroblasts comprised a median of 22% of total fibroblasts in RA compared to 8% in OA (OR=3 (95% CI 1.33–6.48), P=0.007). By contrast, CD34^−^THY1^−^ cells were less abundant in RA at 15% compared to 48% in OA (OR=0.48 (95% CI 0.23–1.03), P=0.06). Within 12 RA samples, the 7 samples that were obtained from swollen joints (according to a rheumatologist's assessment) had different subpopulation proportions than 5 samples from non-swollen joints (**Fig. S5**). The swollen joints had fewer CD34^−^THY1^−^ (P=0.02), more CD34^−^THY1^+^ (P=0.09), and more CD34^+^ fibroblasts (P=0.01). Notably, most cells in the expanded CD34^−^THY1^+^ population in RA also were positive for surface protein levels of CDH11 (median 84%), and CDH11 was also expressed on the other fibroblast subsets including CD34^−^THY1^−^ (median 64%) and CD34+ (median 94%) cells. These results suggest that synovial fibroblast subpopulation proportions are closely related to disease type and activity.

We note that all OA samples were taken from the knee, while RA samples included those from the knee (n=8) as well as other smaller joints (n=8) (**Table S1**). To confirm that the altered proportion of fibroblast subsets in RA reflects the level of tissue inflammation, rather than joint location of origin, we collected independent RA synovial tissue biopsies from only knee joints and examined the proportion of fibroblast subsets and infiltrated leukocytes by flow cytometry (**Table S2**). We selected only samples with synovial hypertrophy on ultrasound images. Similar to our RA versus OA comparison, we observed that the proportion of CD34^−^THY1^+^ fibroblasts is positively correlated with the proportion of infiltrated leukocytes by flow cytometry (**Fig. 4A**). In addition, the proportion of CD34^−^THY1^+^ fibroblasts correlated with both histological synovitis and synovial hypertrophy assessed by ultrasound (**Fig. 4B, C**). These results indicate that the altered proportion of fibroblast subsets in RA reflects tissue inflammation at both the molecular and clinical level. In contrast, the correlation between the proportion of CD34− THY1+ fibroblasts and disease duration was not observed, suggesting that the altered proportion of fibroblast subsets is not a secondary effect of chronic tissue damage (**Fig. 4D**).

**Fig. 4.**
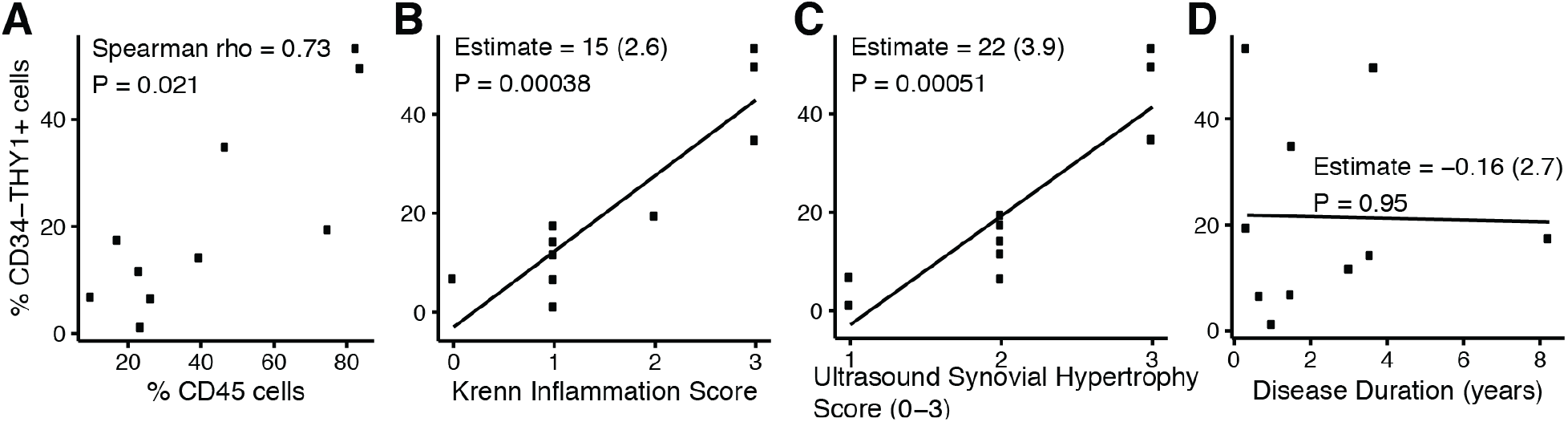
Knee biopsies revealed the correlation between the proportion of fibroblast subsets and synovial inflammation. (**A**) The Spearman correlation between the proportion of CD34^−^THY1^+^ fibroblasts in total fibroblasts and the proportion of CD45+ cells in total live cells in synovial tissue biopsies from RA knee joints. (**B, C, D**) The linear correlation coefficient between the proportion of CD34^−^THY1^+^ fibroblasts and Krenn inflammation score (B), ultrasound synovial hypertrophy score (C), and disease duration (years) (D). Proportion of cells were evaluated by flow cytometry.

### Proportion of fibroblast subsets is associated with accumulation of lymphocytes

We employed mass cytometry to simultaneously assess abundances of fibroblast subsets and infiltrated immune cells in synovial tissues obtained from 18 RA patients with a variety of clinical characteristics (**Table S3**). This enabled us to analyze how the abundances of fibroblast subsets correlated with abundances of infiltrated immune cells in the synovial tissue. The ratio of the number of CD34^−^THY1^+^ to CD34^−^THY1^−^ fibroblasts is positively correlated with the proportion of B cells (P=0.02) and shows a similar trend with the proportion of CD4^+^ T cells (P=0.06) in the synovial tissue (**Fig. S6**). Correlations with other cellular proportions (CD8^+^ T cells, CD14^+^ macrophages) were not observed (**Fig. S6**). These results suggest that increased abundance of CD34^−^ THY1^+^ fibroblasts is associated with accumulation of certain types of lymphocytes within synovial tissue.

### Fibroblast subsets are functionally different

We used gene expression profiles as proxies for molecular functions to investigate functional differences between fibroblast subsets. Gene set enrichment analysis prompted us to investigate how the expression patterns of cytokines are different between fibroblast subsets (**Fig. 5A**) (*13, 14*). Each fibroblast subset expresses distinct types of cytokines and matrix metalloproteinases (MMPs) (**Fig. 5B**). CD34^−^ THY1^+^ fibroblasts express *TNFSF11* (*RANKL*), and lowly express *TNFRSF11B* (*OPG*), a decoy receptor for *TNFSF11* (**Fig. 5B**). The increase of CD34^−^THY1^+^ fibroblasts in RA may explain the increased bone destruction and accumulation of lymphocytes in RA synovial tissue, since RANKL is essential for osteoclastogenesis and is involved in T cell trafficking in autoimmune inflammation (*15, 16*).

**Fig. 5.**
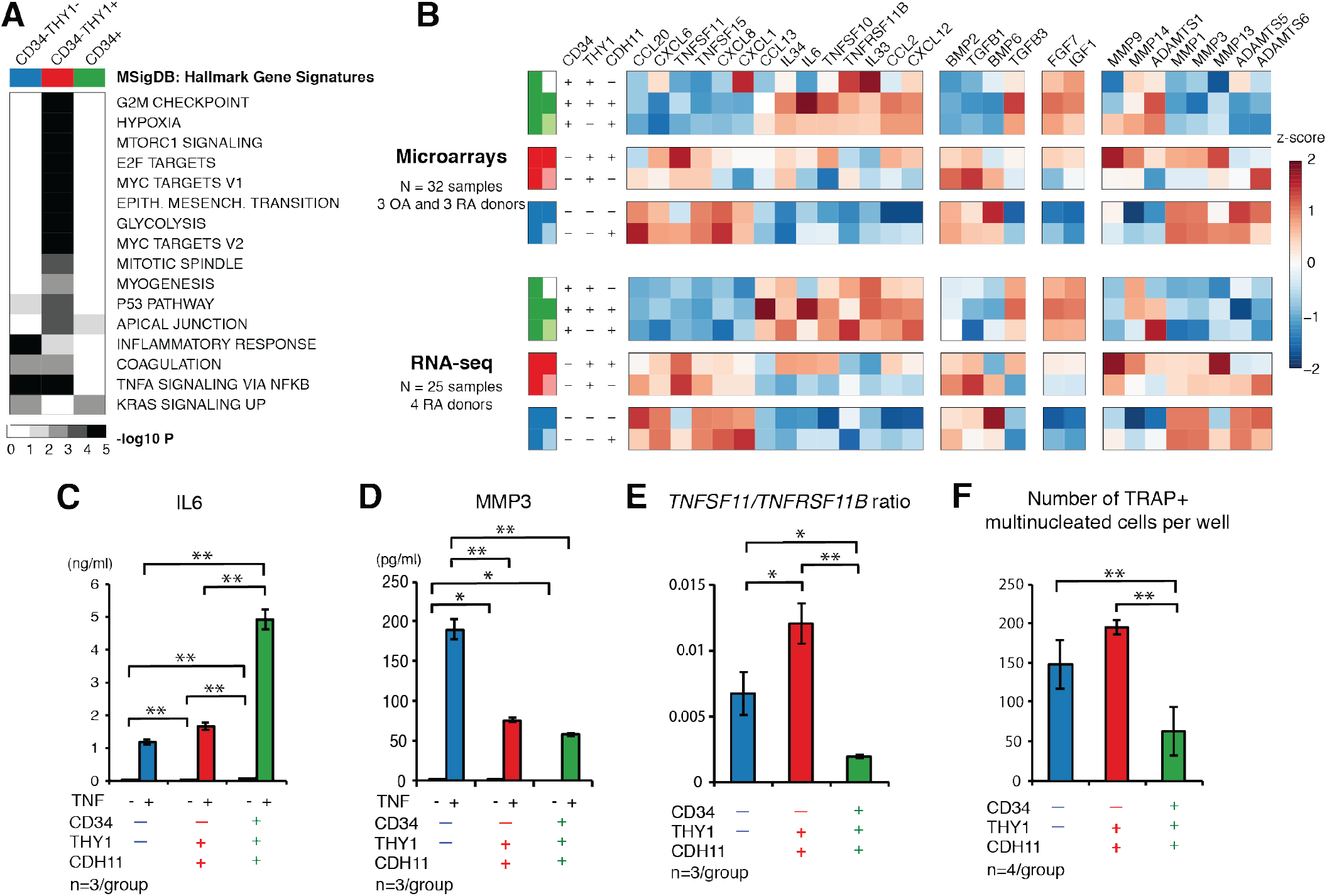
Gene expression profiles in freshly isolated cells and functional assays of cultured cells indicate functional differences between fibroblast subsets. (**A**) Gene set enrichment analysis with RNA-seq samples and MSigDB Hallmark Gene Signatures. Terms with <5% FDR are displayed. (**B**) Expression of selected genes. (**C, D**) Protein expression of IL6 and MMP3 in the supernatant of expanded fibroblast subsets in the presence or absence of 1 ng/ml TNFα. (**E**) The ratio of *TNFSF11* (*RANKL*) and *TNFRSF11B* (*OPG*) in mRNA level in expanded fibroblast subsets. (**F**) Co-culture of osteoclast progenitors with fibroblast subsets in osteoclastogenic conditions. Bars show means and error bars show standard deviations. One-way ANOVA, Tukey post-hoc: * P < 0.05; ** P < 0.01.

CD34^−^THY1^−^ fibroblasts express high levels of *CXCL1* and *CXCL8*, essential for neutrophil recruitment and activation (**Fig. 5B**)(*17*). They also express *BMP6*, known to promote osteoblastic bone formation (**Fig. 5B**)(*18*). Depletion of CD34^−^THY1^−^ fibroblasts may explain the decreased bone formation activity in RA. CD34^+^ fibroblasts express a variety of cytokines including *IL6, CXCL12* and *CCL2*, crucial cytokines that cause tissue inflammation (**Fig. 5B**)(*2, 4*).

To validate our transcriptional analysis, we expanded sorted fibroblast subsets *in vitro* and analyzed their protein expression by ELISA and mRNA expression by qPCR. Consistent with microarray and RNA-seq data in freshly isolated cells, CD34^−^THY1^−^ fibroblasts express high levels of MMP1 and MMP3 in response to TNF stimulation; whereas, CD34^+^ fibroblasts express high levels of IL6 and CXCL12 in response to TNF stimulation (**Fig. 5C, D** and **Fig. S7**). CD34^−^THY1^+^ fibroblasts express high levels of *TNFSF11* (*RANKL*) and low levels of *TNFRSF11B* (*OPG*) relative to CD34^−^THY1^−^ or CD34^+^ fibroblasts (**Fig. 5E and Fig. S7**).

We hypothesized that the differential expression of *TNFSF11* and *TNFRSF11B* between freshly isolated fibroblast subsets would affect their ability to promote osteoclastogenesis, an important effector function for fibroblasts. Indeed, when we cocultured fibroblast subsets with monocytes in osteoclastogenic conditions *in vitro*, we observed that the number of osteoclastic cells developed from monocytes was greater in the co-culture with CD34^−^THY1^−^ fibroblasts or CD34^−^THY1^+^ fibroblasts compared to CD34^+^ fibroblasts (**Fig. 5F**). This suggests that fibroblast subsets differ in their ability to promote osteoclastogenesis and that this function can be detected during short term *in vitro* culture.

## Discussion

This study identifies subsets of fibroblasts in fresh human synovium, including a distinct subset of PDPN^+^CD34^−^THY1^+^ fibroblasts that is expanded in RA and may be pathogenic. These cells are enriched around blood vessels in RA synovium, and their expression profile reveals potential pathogenic roles in lymphocyte recruitment and osteoclastogenesis. We note that almost all of these cells are positive for CDH11, which we have previously shown to be associated with pathological behavior of fibroblasts in *in vitro* studies and RA mouse models (*3*).

Expansion of fibroblasts is a dynamic component of RA synovitis. An increase in synovial lining fibroblasts was noted previously to correlate with disease activity score in 28 joints (DAS28), disease duration, and the level of macrophage infiltration (*19*). Expansion of synovial sublining fibroblasts is also observed in RA, but no previous literature reports significant correlation between sublining fibroblasts and other clinical or pathological findings, except for a negative correlation with DAS28 in one study (*19*). Here, by separating fibroblasts into subsets based on the expression patterns of multiple markers, we found that the increase of CD34^−^THY1^+^ fibroblasts around blood vessels in the sublining area is a dominant change in fibroblasts in RA synovium. Moreover, this expansion distinguishes RA from OA, reflects RA disease activity, and correlates with lymphocyte infiltration in the synovium. Previous studies have shown that mesenchymal CDH11 determines adhesion between fibroblasts, increases their migration and invasion and synergizes in the activation of fibroblasts to produce MMPs, cytokines, and chemokines (*3, 20, 21*). Since a large majority of the expanded fibroblast population expresses this cadherin, it also may contribute to their pathologic behavior in RA (**Fig. 3E**).

There is a real need for strategies to define fibroblast heterogeneity and pathogenic fibroblast populations in order to understand the complex nature of tissue pathology and the role of tissue resident cells in end organ damage. Our approach involves an integrative analysis of cell surface markers, bulk transcriptomes, single-cell transcriptomics, and histological imaging of human tissues that identified a disease-related fibroblast subpopulation that may ultimately serve as a specific target for therapy. We anticipate that such an approach may serve as a template for future studies to identify pathogenic subsets of tissue cells in other human diseases.

## Materials and Methods

### Patient recruitment and isolation of synovial cells

We obtained synovial tissue from joint replacement surgeries, synovectomy surgeries or synovial biopsies of OA or RA patients with appropriate informed consent as required. The study protocols are Institutional Review Board approved at Partners HealthCare, Hospital for Special Surgery, and the University of Birmingham Local Ethical Review Committee. We modified previously described protocols to isolate synovial cells (*22, 23*). Briefly, we obtained tissue immediately after the surgeries. We removed bone and adipose tissues with scissors. We cut synovial tissues into small pieces, and then subjected these pieces to enzymatic digestion. For microarray analysis, low input bulk RNA-seq and *in vitro* assays, we digested tissues with 4 mg/ml collagenase type 4 (Worthington, NJ), 0.8 mg/ml DispaseII, 0.1 mg/ml DNaseI (Roche) in DMEM at 37°C. After 15 minutes, we collected the supernatant and replaced with fresh enzyme mix. We repeated these procedures every 15 minutes for total 1 hour. For the analysis of synovial biopsy samples, we digested the tissues with 0.05 mg/ml Liberase TM (Roche) and 0.04 mg/ml DNaseI at 37°C for 30 minutes. For mass cytometry analysis, we digested tissues with 0.2 mg/ml Liberase TL (Roche), 0.1 mg/ml DNaseI in RPMI at 37°C for 20 minutes to minimize the cleavage of surface markers of lymphocytes during the enzymatic digestion. The same protocol was used to isolate synovial cells for single cell RNA-seq. After lysing red blood cells with ACK-lysing buffer, we stained cells with antibodies, and sorted by FACSAria Fusion (BD) with 100 μm nozzle at 20 psi. For the analysis with microarray and low input RNA-seq, the cells were sorted into 2% FBS HBS+ buffer, spun down, and lysed with TRIzol (Invitrogen). We extracted RNA and cleaned up by RNeasy micro kit (QIAGEN) with DNaseI treatment. For single cell RNA-seq, the cells were stained with antibodies and directly sorted into 5 μl of TCL buffer (QIAGEN) with 1% β-mercaptoethanol (SIGMA) in 96 well plates. For the analysis with mass cytometry, we froze isolated synovial cells in CryoStor CS10 (STEMCELL Technologies, Vancouver, Canada) for batched analysis.

### Fibroblast markers

#### Selection of markers

We selected fibroblast markers for separation of putative fibroblast subsets on the basis of published literatures. PDPN is known to be a marker for fibroblastic reticular cells (FRCs) in lymph nodes and is highly expressed in fibroblasts in synovial tissues, especially in lining fibroblasts (*24, 25*). CDH11 is also highly expressed in FRCs and lining fibroblasts in RA synovial tissue (*24, 26*). THY1 has been widely used as a fibroblast marker in cultured cells, and is highly expressed in sublining fibroblasts in synovial tissue (*25, 27*). Expression of CD34 has been documented in fibroblasts and endothelial cells (*28, 29*). CD146 has been used as a marker for pericytes (*29*).

#### Antibodies and reagents

The following antibodies and reagents were used for the analysis of synovial cells with flow cytometry and cell sorting: anti-CD45-APC-H7 (2D1, BD Pharmingen), anti-CD235a-APC-Alexa Fluor750 (11E4B-7-6(KC16), Beckman Coulter), anti-CD31-PE-Cyanine7 (WM-59, eBioscience), anti-CD146-BV450 (P1H12, BD Horizon), anti-CD34-PE (4H11, eBioscience), anti-Podoplanin-PerCP-eFluor710 (NZ-1.3, eBioscience), anti-THY1-FITC (5E10, BD Pharmingen), anti-Cadherin11-Biotin (23C6), human TruStain FcX (Biolegend), Streptavidin-APC (Jackson ImmunoResearch), Live/Dead fixable aqua dead cell stain kits (Molecular probes).

For immunofluorescence staining of synovial tissue, following antibodies and reagents were used: anti-CD45 (135-4C5, Abd serotec), anti-CD34 (EP373Y, Abcam), anti-Podoplanin (NZ-1.3, eBioscience), anti-THY1 (F15-42-1, Merck Millipore), anti-mouse IgG1-FITC (Southern Biotech), anti-mouse IgG2b-Alexa Fluor 647 (Life Technologies), anti-rat IgG-Alexa Fluor 594 (Life Technologies), anti-rabbit IgG-Alexa Fluor 546 (Life Technologies), and Hoechst 33258 (Life Technologies).

### Transcriptional analysis

#### Gene expression microarrays

We evaluated the integrity of RNA with Bioanalyzer or by Tapestation (Agilent). We used only RNA with more than RIN score of 7. We prepared cDNA from 38.1 μg total RNA using Ovation Pico WTA (NuGEN), followed by labeling 5 μg cDNA using Biotin Module (Nugen). We assayed gene expression using the GeneChip Human 2.0 ST microarray (Affymetrix). We normalized expression by Robust Multiarray Averaging (RMA). We identified and removed two outlier arrays by principal components analysis (PCA). We assigned probes to Entrez Gene IDs using Ensembl BioMart on March 17, 2015. In the instance where there were multiple probesets assigned to a single gene, we averaged them to obtain a single gene value.

#### RNA-seq library preparation and sequencing

We used 1,250 cells per sample for library preparation. We prepared sequencing libraries using the Smart-Seq2 protocol. We pooled and sequenced libraries were pooled and sequenced with the Illumina HiSeq 2500 to a depth of 8–14M reads per library.

#### Single-cell RNA-seq sequencing

We assayed 384 fibroblasts from 4 donors, 2 with OA and 2 with RA. For each donor, we collected fresh synovial tissue, isolated synovial cells by enzymatic digestion, and stained with antibodies against CD45, CD235a, CD31, CD146, PDPN, CD34, THY1 and CDH11. We sorted 96 single CD45-CD235a-CD31-PDPN+ cells by FACSAria Fusion (BD), and assayed mRNA expression with the Smart-Seq2 protocol (*30*). Single-cell libraries were also prepared with the same protocol, and we aimed to sequence to a depth of 200K-12M reads per library (*31*). On average, we sequenced 5.4M fragments and detected 8,153 genes per cell with at least 1 TPM. We discarded 47 cells (12%) with fewer than 5,000 genes detected from further analysis.

#### RNA-seq gene expression quantification

We quantified cDNAs on canonical chromosomes in Ensembl release 83 with Kallisto v0.42.4 (*32*) in transcripts per million (TPM) and summed to get gene-level expression values. We quantified gene expression the same way for bulk and single cell RNA-seq samples. For differential expression analysis, hierarchical clustering, and principal components analysis (PCA), we log (base 2) transformed TPM values.

#### Lineage marker analysis of microarrays, bulk RNA-seq, and single-cell RNA-seq

We selected lineage markers for fibroblast, endothelial, and hematopoietic cells (*33*) and checked their expression levels to confirm that our samples are from the fibroblast lineage (**Fig. S1**).

#### Differential expression analysis with microarrays

We used the R package limma to assess differential expression analysis on RMA normalized expression values (*34*) Before performing differential expression analysis, donor-specific variation was regressed out by obtaining the residuals from linear models. Each gene was modeled as a linear combination of donor-specific effects, and the residuals from these models were used for differential expression analysis, hierarchical clustering, and PCA. We expected differences in gene expression between RA and OA. However, we lacked power to see these differences within fibroblast subsets.

#### Differential expression analysis with RNA-seq

RNA-seq data was analyzed the same way as the microarray data, starting with log2 transformed TPM expression values. Donor-specific variation was similarly regressed out before differential expression analysis, hierarchical clustering, and PCA.

#### Gene set enrichment analysis

Terms from MSigDB Hallmark Gene Signatures were used for enrichment analysis with Gene Set Enrichment Analysis (*35*). We tested gene sets for enrichment with differential expression (DE) between the 3 major subsets in order to assess how they might differ from each other in terms of molecular pathways. MSigDB hallmark pathways enriched with DE signal for CD34-THY1+ population, expanded in RA joints, include “epithelial to mesenchymal transition”, “hypoxia”, and “glycolysis” (*36*).

#### Principal components analysis (PCA)

After gene selection by ANOVA or differential expression analysis, we scaled each sample and then scaled each gene across the samples to obtain a specificity of the gene to each sample. Next, we used the prcomp function in R to perform PCA with centered and scaled log2 expression values.

#### Linear discriminant analysis (LDA)

We checked if transcriptional profiles of single cells are similar to profiles of the 3 major subsets we defined by bulk transcriptomics. First, we selected 1,171 genes with significant (5% FDR) differential expression between any pair of the 3 subsets in bulk RNA-seq data. We selected a subset of 968 genes with high expression in single cells (mean log2(TPM) > log2(10)). We used the bulk RNA-seq data to train a linear discriminant analysis (LDA) model with these genes, and then classified each single cell's expression profile to predict each single cell’s identity. The confidence of each classification is represented by a posterior probability.

### Histological analysis

RA synovial tissues were obtained by biopsies from RA patients in the BEACON Birmingham early arthritis cohort, which is an early arthritis cohort recruiting patients with new onset arthritis prior to treatment with disease-modifying antirheumatic drugs. Synovial tissues for staining were frozen in OCT compound. Sections were made in 6 μm thickness, fixed with acetone, and frozen prior to use. Slides were rehydrated in PBS, blocked with 10% normal goat serum in PBS for 10 minutes, and then incubated with primary antibodies, followed by secondary antibodies. Slides were mounted using ProLong Diamond (Life Technologies), and imaged using a Zeiss LSM 780 confocal microscope. Images were processed using Zen Black and Zen lite (Zeiss). Representative images were shown. Synovial tissues for histological analysis and haematoxylin and eosin staining were fixed in formaldehyde then mounted in paraffin, sectioned and stained by the Hospital Pathology service.

### Histological evaluation of inflammatory infiltrate

Hematoxylin and eosin-stained sections of knee synovial biopsy samples were examined histologically for the severity of inflammatory infiltrate using the inflammatory component of the Krenn synovitis score (*37*). Inflammatory infiltrates were graded from 0–3 (0=no inflammatory infiltrate, 1=few mostly perivascular situated lymphocytes or plasma cells, 2=numerous lymphocytes or plasma cells sometimes forming follicle-like aggregates, and 3= dense band-like inflammatory infiltrate or numerous large follicle-like aggregates). The tissues were graded in a blinded manner by two trained individuals then consensus agreed.

### Clinical evaluation of synovitis by ultrasound

The joint to be biopsied was assessed using ultrasound immediately prior to the procedure using a Siemens Acuson Antares scanner (Siemens PLC, Bracknell, UK) and multifrequency (5–13MHz) linear array transducers. Synovitis and power Doppler (PD) positivity were defined using consensus OMERACT definitions (*38*). Greyscale synovial hypertrophy and Power Doppler ultrasound variables were graded on 0–3 semi-quantitative scales as previously reported (*39*).

### Functional analysis

#### Cell culture

We sorted CD34−THY1− fibroblasts, CD34−THY1+CDH11+ fibroblasts, and CD34+THY1+CDH11+ fibroblasts, and cultured them in Dulbecco’s modified Eagle’s medium (DMEM) supplemented with 10% fetal bovine serum (FBS; Gemini), 2 mM L-glutamine, antibiotics (penicillin and streptomycin), and essential and nonessential amino acids (Life Technologies). The cells were expanded for 3-20 days for assays *in vitro*. The cells with one or two passages were used. The cells were cultured in the presence or absence of 1 ng/ml of TNF-α (R and D) for 24 hours for ELISA of IL-6, CXCL12, MMP-1, MMP-3 and MMP-14, or for 72 hours for qPCR of *TNFSF11* and *TNFRSF11B*.

#### ELISA

The levels of IL-6, CXCL12, MMP-1, MMP-3 and TNFRSF11B in the supernatant or the levels of MMP-14 in the cell lysate were evaluated by ELISA kit as described in manufacturer's instructions (Duo Set, R and D).

#### Quantitative Real-Time PCR (qPCR)

cDNA was synthesized with QuantiTect Reverse Transcription kit (QIAGEN). qPCR was performed with Brilliant III Ultra-Fast SYBR Green qPCR master mix (Agilent Technologies) on a Mx3000 (Stratagene). Following primers were used; *TNFSF11*, forward; 5’-GGA GAG GAA ATC AGC ATC GAG, reverse; 5’-CCA AAC ATC CAG GAA ATA CAT AAC AC, *TNFRSF11B*, forward; 5’-CAA CAC AGC TCA CAA GAA CAG, reverse, 5’-GAA GGT GAG GTT AGC ATG TCC, *GAPDH* forward; 5’-AAT CCC ATC ACC ATC TTC CAG, reverse; 5’-AAA TGA GCC CCA GCC TTC.

#### Osteoclastogenesis assay

Osteoclast progenitors were prepared by culturing PBMC in the presence of 20 ng/ml of M-CSF (PeproTech) in DMEM supplemented with 10% FBS (GE Healthcare), 2 mM L-glutamine and antibiotics for 5–6 days. Expanded fibroblasts were seeded at 5,000 cells/well in 96 well plate. On the next day, osteoclast progenitors were added at 5,000 cells/well, and were co-cultures with fibroblasts in the presence of 20 ng/ml M-CSF and 5–20 ng/ml RANKL (PeproTech). The media was replaced every 2 days. After 6 days of the co-culture, the cells were fixed by 4% PFA. After TRAP staining, TRAP positive multinucleated cells were counted as osteoclasts.

#### Mass cytometry

After thawing cryopreserved synovial cells, the cells were first stained with cisplatin for the assessment of viability, and then stained with antibody cocktails as described previously (*40*). The cells were fixed and permed with intracellular fixation and permeabilization buffer (eBioscience), stained for intracellular markers, and re-fixed with formalin (Sigma). The stained cells were analyzed with Helios (Fluidigm, CA, USA). Data were normalized using EQ^TM^ Four Element Calibration Beads (Fluidigm), and were analyzed with FlowJo 10.1 (FlowJo, OR, USA).

#### Statistical analysis

The statistical analysis for each experiment was described in sections of results, figure legends, and supplementary materials.

## Acknowledgments

We thank the BWH Human Immunology Center Flow Cytometry Core for cell sorting assistance, Partners Personalized Medicine Translational Genomics Core for microarray analysis and Broad Technology Labs for RNA-seq.

## Author Contributions

F.M. conceived the project, performed experiments, analyzed data, and wrote the manuscript. K.S. analyzed microarray and RNA-seq data, and wrote the manuscript. J.M. and A.F. performed histological analysis. K.W., D.A.R., S.K.C., H.N.N., E.H.N., J.D.T performed experiments. B.E.E., P.E.B., J.W., B.P.S., G.K., S.M.G., V.P.B., L.B.I., P.A.N participated in patient recruitment, sample acquisition. A.F. and C.D.B. participated in patient recruitment and sample acquisition, and contributed to the review and interpretation of the data. L.T.D. participated in patient recruitment, sample acquisition and review of the data. J.A.L. developed reagents and assisted with mass cytometry. N.H. assisted with RNA-seq. S.R. co-supervised the project, analyzed data, and co-wrote the manuscript. M.B.B. conceived the project, supervised the work, analyzed data, and co-wrote the manuscript.

## Data and materials availability

Microarray, RNA-seq, flow cytometry, and mass cytometry data will be made publicly available at GEO.

## Code availability

R source code for the analysis is available upon request.

## Supplementary Materials

Fig. S1. Heatmap of expression data for genes characteristic of cell types in synovial tissues.

Fig. S2. Concordance of differential expression between microarrays and low-input RNA-seq.

Fig. S3. Assessing quality of RNA-seq data by visualizing the distributions of number of cDNA fragments assigned to mRNA transcripts and the number of genes detected with at least 1 transcript per million (TPM).

Fig. S4. Agreement between protein gating and LDA classification based on single cell RNA-seq.

Fig. S5. Proportion of fibroblast subsets in swollen and non-swollen joints in RA patients.

Fig. S6. Correlations between the ratio of CD34^−^THY1^+^ to CD34^−^THY1^−^ synovial fibroblasts and other cellular and clinical parameters.

Fig. S7. The expression of cytokines and MMPs *in vitro*.

Table S1. Clinical characteristics of evaluated patients with flow cytometry.

Table S2. Clinical characteristics of patients who donated synovial biopsy samples from knee joint.

Table S3. Clinical characteristics of evaluated patients with mass cytometry.

